# Anellovirus Structure Reveals a Mechanism for Immune Evasion

**DOI:** 10.1101/2022.07.01.498313

**Authors:** Shu-hao Liou, Noah Cohen, Yue Zhang, Nidhi Mukund Acharekar, Hillary Rodgers, Saadman Islam, Lynn Zeheb, Jared Pitts, Cesar Arze, Harish Swaminathan, Nathan Yozwiak, Tuyen Ong, Roger J. Hajjar, Yong Chang, Kurt A. Swanson, Simon Delagrave

## Abstract

The significant impact of the human virome on human physiology is beginning to emerge thanks to modern sequencing methods and bioinformatic tools^1^. Anelloviruses, the principal constituent of the commensal human virome, are universally acquired in infancy and found throughout the body^2,3,4^. Since the discovery of the original torque teno virus in 1997^5^, three genera of the *Anelloviridae* family, each extremely diverse genetically, have been found in humans. These viruses elicit weak immune responses that permit multiple strains to co-exist and persist for years in a typical individual^6^. However, because they do not cause disease^7^ and due to the lack of an *in vitro* culture system, anelloviruses remain poorly understood^8,9^. Basic features of the virus, such as the identity of its structural protein, have been unclear until now. Here, we describe the first structure of an anellovirus particle, which includes a jelly roll domain that forms a 60-mer icosahedral particle core from which spike domains extend to form a salient part of the particle surface. The spike domains come together around the 5-fold symmetry axes to form crown-like features. Relatively conserved patches of amino acids are near the base of the spike domain while a hypervariable region is at the apex. We propose that this structure renders the particle less susceptible to antibody neutralization by hiding vulnerable conserved epitopes while exposing highly diverse epitopes as immunological decoys, thereby contributing to the immune evasion properties of anelloviruses. This would contrast with viruses such as beak and feather disease virus, canine parvovirus or adeno-associated virus which lack such pronounced surface features. These results shed light on the structure of anelloviruses and provide a framework to understand their interactions with the immune system.

## Main Text

Anelloviruses comprise a single-stranded, negative-sense circular DNA genome ranging from ∼2.0-3.9 kilobases (kb). Among the three genera of human anelloviruses, the *Alphatorquevirus* (torque teno virus; TTV) genome is ∼3.8 kb, the *Gammatorquevirus* (torque teno midi virus; TTMDV) genome is ∼3.2 kb, and the *Betatorquevirus* (torque teno mini virus; TTMV) genome is ∼2.9 kb. Through alternative splicing, six or seven different open reading frames (ORFs) are encoded by these genomes, and the longest one is known as ORF1^10^. A distinguishing feature of ORF1 is the presence at the N-terminus of several arginine residues and some lysines which are consistently observed in anelloviruses of all genera. This arginine-rich motif (ARM) is seen in several viral families and is thought to play a role both in nuclear localization signaling and viral genome binding ^11^. ARM-containing viruses typically contain a jelly roll (JR) domain characteristic of viral capsid proteins^12^. Therefore, we hypothesized that the ORF1 protein also contains a JR domain. However, no experimental evidence was available to confirm the presence of such a domain in the anellovirus ORF1. Indeed, a few failed attempts at expressing full-length ORF1 have been reported by others^10^, limiting characterization of this protein.

### Anellovirus LY1 ORF1 fragment structure

Our initial efforts to study the structure of a virus-like particle (VLP) derived from a Betatorquevirus isolate called LY1^13^, were performed with the full-length ORF1 (residues 1-672) expressed in insect cells (Sf9). Full-length ORF1 assembled into particles ∼32 nm in diameter (Extended Data Fig. 1A), similar to previously reported estimates of anellovirus size^14^. However, full-length ORF1 particles lacked the homogeneous symmetry expected of viral particles. Genome packaging by ARM-containing viruses is believed to occur when the positive arginine residues bind a negatively charged genome, overcoming electrostatic repulsion of the ARM to permit ORF1 assembly. We therefore designed LY1 ΔARM, an ORF1 construct wherein residues 2-45 are deleted (Fig. 1A,B and Extended Data Figure 1B) to facilitate more efficient ORF1 assembly in the absence of viral DNA. LY1 ΔARM expression was confirmed using a rabbit polyclonal antibody raised against a spike P1 domain peptide (residues 485-502; Fig. 1B). We observed a band consistent with LY1 ΔARM by western blot above the 62-kDa marker (expected mass of 73.3 kDa; Fig. 1C). After cell lysis and purification we observed the ORF1 band was below the 62-kDa marker. The difference in apparent molecular weight of LY1 ΔARM fragment after initial expression and after purification suggested proteolysis. To identify the site of proteolysis, we generated polyclonal antibodies to peptides from JR domain at the extreme N-terminus (residues 46-58) and C-terminal domain at the C-terminus of LY1 ΔARM (residues 635-672; Fig. 1B) and confirmed the presence of the N-terminal peptide of LY1 ΔARM fragment and the absence of the C-terminal peptide (Fig. 1C).

**Fig. 1.**
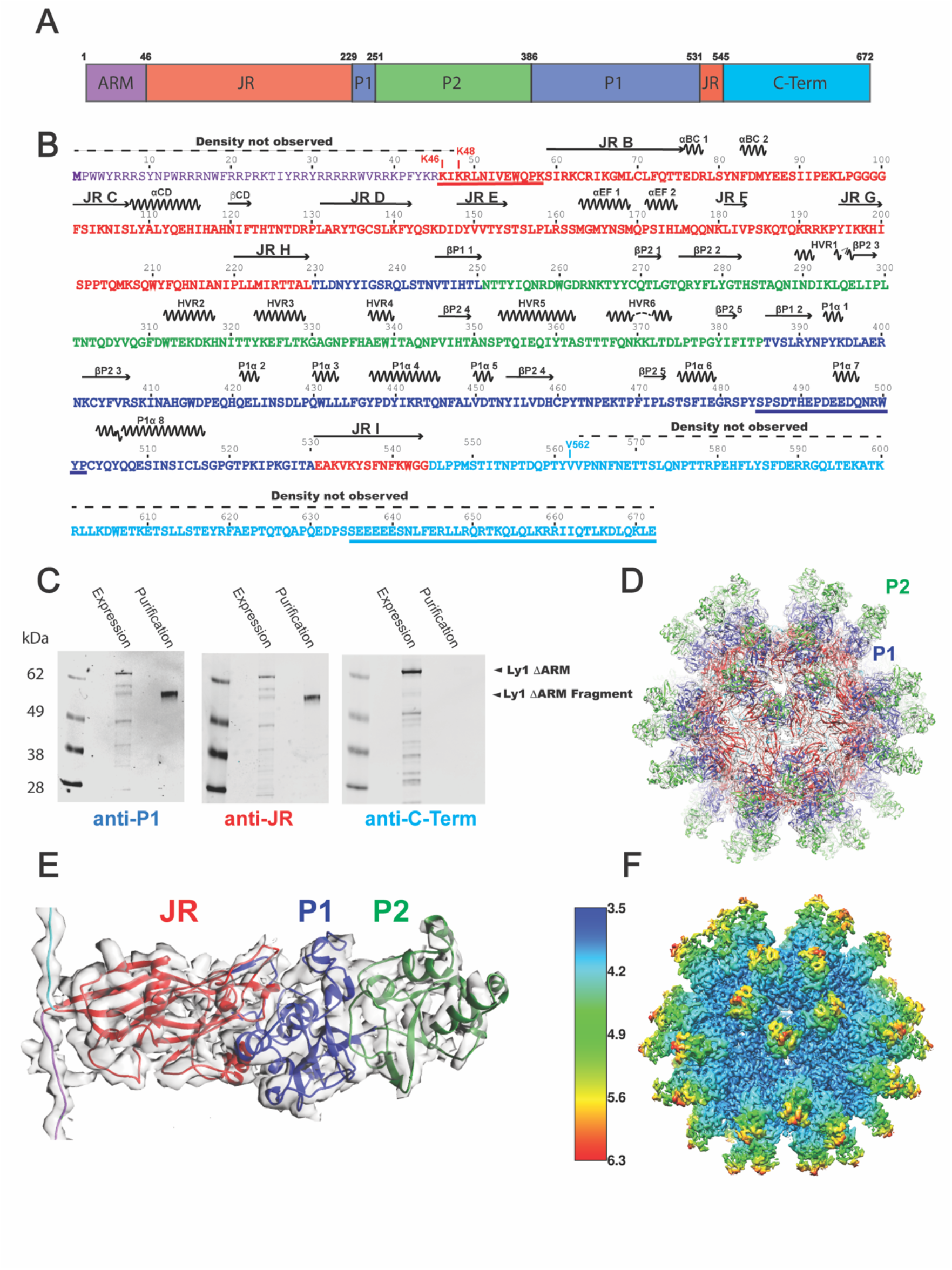
Anellovirus LY1 ΔARM construct design and proteolysis. **A.**A schematic representation of full-length LY1 ORF1 is shown as a cartoon labeled and colored by domains. The arginine-rich motif (ARM) is shown in purple, the jelly roll (JR) domain is shown in red, the spike P1 domain is shown in blue, the spike P2 domain is shown in green, and the C-terminal domain is shown in cyan. Residue numbers beginning each domain and the C-terminus are indicated above. **B**. The sequence of full-length LY1 ORF1 colored as in A with residue numbers indicated above. In bold are residues included in the LY1 ΔARM construct including the initial residue, K46, which is labeled. A dashed line above the sequence indicates residues not observed in the electron density. Secondary structure elements are indicated above with β-strands as arrows and α-helices as zig-zag lines. The JR β-strands are labeled B-I per convention while additional secondary structures are numbered by their domain. Three peptides used to generate polyclonal antibodies are underlined. **C**. Western blot analysis of LY1 ΔARM after expression (Expression) and after purification (Purification). A molecular weight marker is labeled to the left of the gels, while arrows on the right indicate the band of LY1 ΔARM before (LY1 ΔARM) and after proteolysis (LY1 ΔARM Fragment). Polyclonal antibodies used to probe the western blots, colored and named by the peptides used to generate them (A), are indicated below. **D**. The 3D reconstruction of LY1 ΔARM VLP electron density and 60-mer VLP molecular structure colored as in A. The spike P1 and P2 domains are labeled. **E**. One ORF1 protomer shown in its electron density with domains labeled and colored as in A. **F**. The electron density of LY1 ΔARM VLP colored by its local resolution. A bar indicates the resolution (unit in angstrom) scale by color. The particle is oriented as in D.

Electron microscopy (EM) analysis of the LY1 ΔARM fragment showed the VLPs formed were more homogeneous and symmetric in morphology relative to full-length ORF1 (Extended Data Fig. 1). The formation of homogeneous VLPs following genetic removal of the N-terminus and proteolysis of the C-terminus has been observed in another JR-containing virus, hepatitis E (HEV)^15^. To determine if the C-terminus of LY1 ORF1 is required for particle formation, we generated VLPs from construct LY1 ORF1 ΔARM ΔC-Term (Δ2-45 and Δ552-672). LY1 ORF1 ΔARM ΔC-Term produced VLPs of similar symmetry to LY1 ORF1 ΔARM (Extended Data Fig. 1C). These results suggest that proteolysis of the ORF1 C-terminus may be a natural part of anellovirus formation. In light of recent evidence that the C-terminal region of ORF1 is the immunodominant region of anelloviruses^16^, its excision from the mature particle would be consistent with the immune evasion properties of anelloviruses^17^. Further exploration of the role of the C-terminal region of ORF1 will help clarify its importance in the anellovirus life cycle.

We determined the structure of the LY1 ΔARM particle using cryo-EM to 3.98 Å resolution (Fig. 1D-F, Extended Data Fig. 2 and 3, Extended Table S1). The anellovirus particle is formed by sixty ORF1 fragments organized in an icosahedral T=1 symmetry (Fig. 1D). Electron density for residues 48-562 is observed (Fig. 1B,E). The resulting mass of the observed LY1 fragment is calculated to be ∼59.8 kDa, consistent with the observed mass by western blot (Fig. 1C). The N-terminal region (residues 46-228) forms part of the canonical 8-β–strand JR domain (β strands named B to H by convention). Unexpectedly, the eighth and final β–strand in the JR (strand I, residues 531-542) is located just prior to the C-terminal domain (Fig. 1A,B). The resulting fold of the ORF1 protomer has residues at the N- and C-termini generating the JR domain at the particle core while the intrastrand residues form the exterior of the particle surface. Intrastrand insertions forming the viral particle exterior can be found in other JR-containing viruses such as adeno-associated virus (AAV) and canine parvovirus (CPV) ^18,19^. However, while the intrastrand insertions for AAV2 (228 residues) and CPV (227 residues) are between G and H β–strands, the 298-residue intrastrand insertion in LY1 is significantly larger and lies between β–strands H and I. The intrastrand region (residues 229-530) extends from the JR domain to form a structure herein referred to as a spike domain. The spike domain is formed by two globular domains: the spike P1 domain (residues 229-250 and 386-530) and the spike P2 domain (residues 251-385; Fig. 1B,E).

### Jelly roll domains

Sixty LY1 JR domains form the core of the virus particle (Fig. 2). The β–strands form β–sheets which are characterized by a C-H-E-F pattern on the core’s exterior and B-I-D-G pattern on the core’s interior. The N-terminus of strand B is oriented to place the ARM on the interior of the core, where it is positioned to bind the viral genome. The observed C-terminal residues (545-562) extend from the C-terminus of β-strand I on the interior of the particle and thread through JR domains on the 2-fold axis to contact the neighboring JR domain (Fig. 2B).

**Fig. 2.**
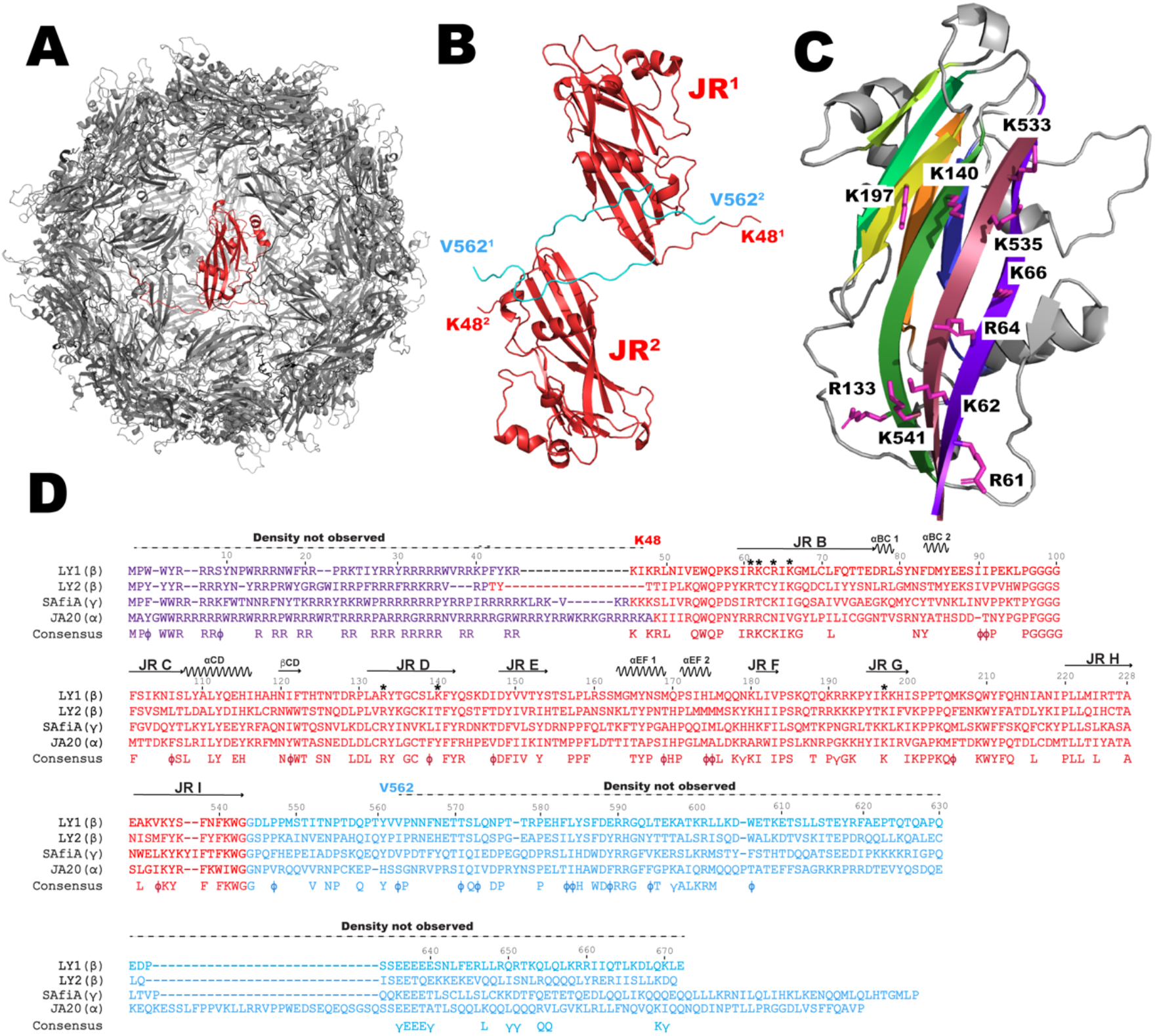
Sixty LY1 jelly roll (JR) domains form the core of anellovirus particles. **A.**The LY1 core structure comprised of 60 JR domains pack in icosahedral symmetry with one domain uniquely colored in red. **B**. Two JR domains are shown in red with the observed C-terminal domain backbone colored in cyan. The JR domains are arbitrarily labeled JR^1^ and JR^2^ with the first (K48) and last (V562) observed residues for each protomer labeled with the corresponding number for clarity. **C**. A single JR domain is oriented to show the β-sheet on the interior of the particle core. Sidechains of basic residues in position to contact with the viral genome are shown and labeled. **D**. The arginine-rich motif, JR, and C-terminal domains of LY1 are aligned with ORF1 sequences from different anellovirus genera (indicated in parentheses). Basic residues of LY1 positioned to potentially contact the viral genome are indicated with asterisks.

In several JR-containing viruses^20^, positively charged residues (arginine and lysine) oriented internally on strands B, I, D, and G are expected to bind the negatively charged viral genome (Fig. 2C). In LY1, basic residues Arg61, Lys62, Arg64, Lys66 (β–strand B), Arg133, Lys140 (β–strand D), Lys197 (β–strand G), Lys533, Lys535, and Lys541 (β–strand I) are all oriented toward the particle interior and are likely responsible, together with the ARM motif, for binding the negatively charged viral genome. Notably, we do not observe any electron density suggesting bound nucleic acid, possibly because the ARM deletion prevented nucleic acid binding or because any host cell nucleic acid encapsidated by the VLP would be too heterogeneous for detection. Alignment of anellovirus ORF1 sequences reveals that several of these putative DNA-binding residues are conserved across species, which supports their role in DNA-binding (Fig. 2D).

### Spike domains

Residues 229 to 530 form the spike domain that extends ∼6 nm from the JR core (Fig. 3). A β-strand (residues 245-250) extending from JR β–strand H is the first component of the spike P1 domain and is N-terminal to the spike P2 domain (residues 251-385). The previously described hypervariable region (HVR) of ORF1 comprises the majority of the spike P2 domain. The remaining residues of the spike domain (residues 386-530) form five additional β–strands and eight helices, which, together with the residues 245-250 strand, fold into the spike P1 domain. The local resolution of P1 is only slightly lower than the JR domain (∼4-4.5 Å), while the resolution of P2 is within 5-6 Å. This is likely a consequence of both being further from the radius of gyration and some flexibility of the HVR residues.

**Fig. 3.**
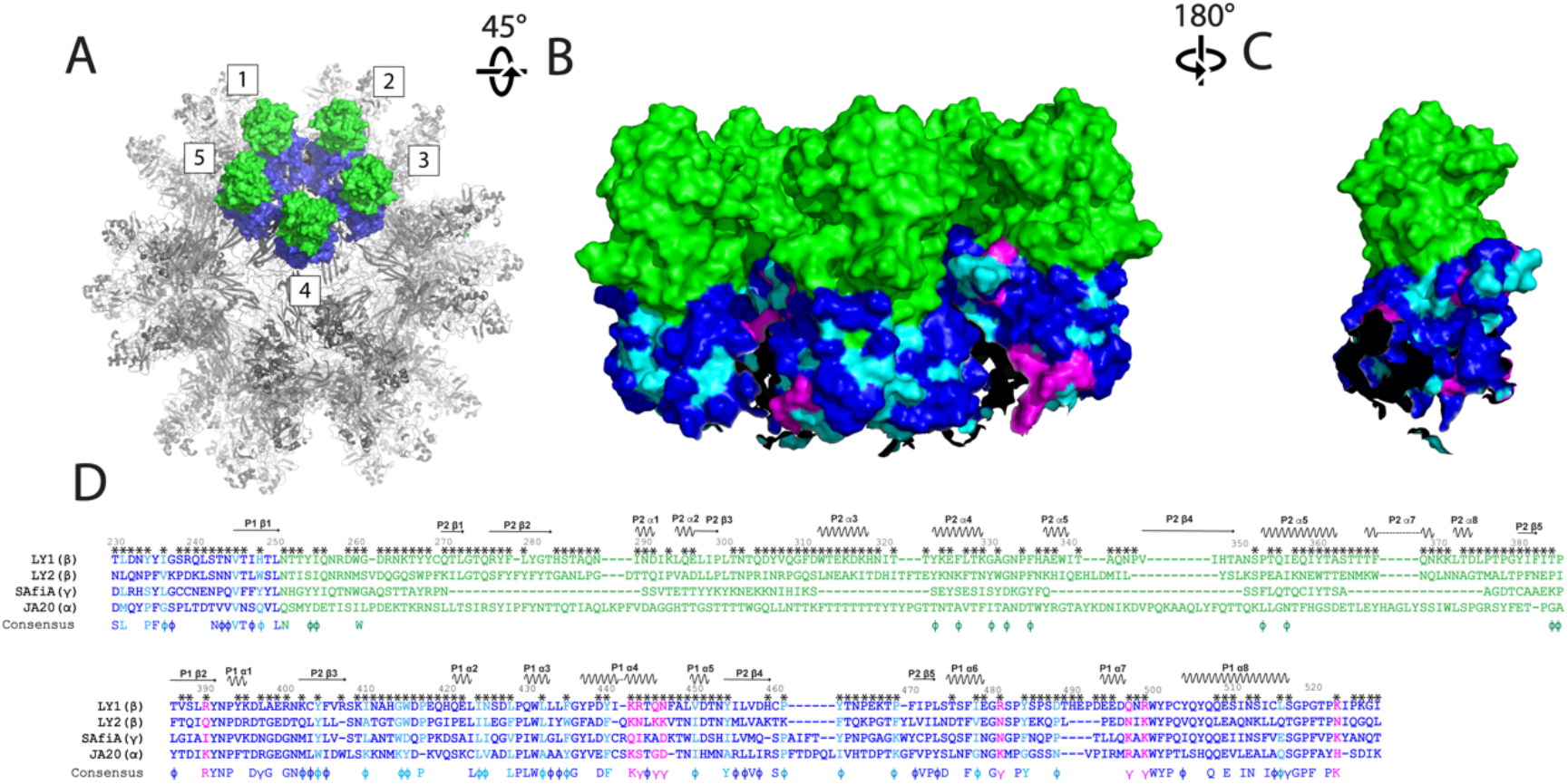
The spike domains extend from the core on the 5-fold axis. **A.** The anellovirus particle structure shown as a surface rendering. The particle is shown in gray with 5 spikes forming a crown structure, numbered for clarity, shown in surface rendering and colored as in Fig. 1. **B**. The exterior of the crown structure shown from the side. Five spike domains are colored as in A. The surface-exposed hydrophobic and hydrophilic residues from the consensus sequence are colored in cyan and magenta, respectively. **C**. The front spike domain from B rotated to view residues on the interior of the crown structure. **D**. The spike domain of LY1 (colored as in Fig. 1 with hydrophobic and hydrophilic residues from the consensus colored as above) is aligned with ORF1 sequences representative of different anellovirus genera (indicated in parentheses). Black asterisks indicate residues that are at least partially surface exposed. Below the alignment is the consensus sequence colored as above.

Neighboring spike domains pack together around the five-fold symmetry axis to form a ringed structure of 5 spike domains henceforth called the crown (Fig. 3A,B). A receptor for anelloviruses has not been identified to date. Given the diverse tropism of different anellovirus strains, it is possible that residues of the spike P2 HVR, which are the most surface-exposed, are involved in viral attachment and infection. However, we hypothesize that the hypervariable sequence of the spike P2 domain serves to aid in immune evasion rather than harboring a receptor-binding motif. If this is the case, a receptor-binding motif on the better conserved spike P1 domain, or even on the surface of the JR core, may exist.

Sequence alignment of the LY1 spike domain with other anelloviruses does not readily identify conserved spike surface residues (Fig. 3B-D, Extended Data Figure 4). In fact, the LY1 residues that are more conserved (at least 50%) between other anelloviruses (e.g., Asp396, Pro429, Trp431, Gly437, Phe477, Pro483, Trp500, Tyr501, Pro502, Gly518 and Pro519) are internal spike P1 residues supporting its globular fold. A few basic residues are at least 50% conserved on the spike P1 surface (e.g., Arg390, Arg481, Arg499, and Arg523) which could contribute to the particle’s ability to bind heparin resin. A few hydrophobic or bulky aromatic (Tyr) residues on spike P1 (e.g., Leu231, Tyr234, Ile236, Val245, His248, Tyr404, Ile410, Gly414, Ile424, Leu428, Leu432, Tyr440, Val450, Tyr454, Pro461, Tyr462, Pro469, Ile478, Tyr484 and Leu516) are at least partially surface-exposed and could represent a conserved receptor binding surface (Fig. 3B-D). If this is the case, then it is attractive to posit that anelloviruses have evolved their novel elongated H-I intrastrand spike domain to sterically hinder antibody binding to their cellular receptor-binding site present on the P1 surface using domain P2 which is able to tolerate highly diverse amino acid substitutions. This would allow diverse anelloviruses to repeatedly infect human hosts with minimal recognition and neutralization by the immune system.

Absent from the LY1 structure is the majority of the C-terminal region (residues 545-672). Observed density for residues 545-562 shows that, were the C-terminal region present on the wild-type virus, the C-terminus would extend from the JR core toward the surface near the 3-fold axis between the spike domain crowns. The C-terminal region is predicted to be helical in nature, and has a glutamate-rich region (residues 636-640 on LY1) which could N-terminally cap a conserved helical motif. To determine if the C-terminal residues of LY1 would form a helical structure, we performed circular dichroism experiments on the C-terminal peptide used to generate the aforementioned antibodies. Indeed, circular dichroism shows the C-terminal region (residues 635-672) is helical in solution suggesting the C-terminal region would form a coiled-coil domain (Extended Data Figure 5) in a leucine-zipper like association of the helices consistent with the observed periodicity of hydrophobic residues. Electron microscopy comparing full-length ORF1 vs ΔARM ΔC-Term ORF1 suggests particle formation is improved with the absence of the C-terminus (Extended Data Figure 1C). Given that anelloviruses have evolved to evade the immune system, and that the C-terminal region is the immunodominant region of ORF1 and may be antagonistic to particle formation, we hypothesize that the helical C-terminal region is required for wild-type particle assembly but somehow modified during viral maturation to permit particle formation.

## Discussion

Despite anelloviruses constituting the majority of the human virome, their capsid structure was unknown and the identity of the capsid protein itself had not been established experimentally^21^. Our determination of the LY1 structure demonstrates that ORF1 encodes the capsid protein and that anelloviruses evolved a novel spike domain that extends around the 5-fold axis to form a crown structure. These crowns are capped with hypervariable P2 regions, which likely inhibits the development of antibodies against the better conserved spike P1 domain via steric hindrance. Analysis of the P1 surface reveals conserved residue patches that may have a receptor binding function.

The structure of the LY1 *Betatorquevirus* can now be used to guide future anellovirus research. In particular, the diversity and immunological stealth of anelloviruses indicates that they could be used to deliver therapeutic genes to cell types not currently addressed by existing vectors, and that they may be less susceptible to pre-existing immunity or to the development of neutralizing antibodies following initial treatment^17^. The availability of a capsid structure will help guide the design of anellovirus-based gene therapy vectors and support the interpretation of upcoming experimental data.

## Methods

### Antibody generation and western blot analysis

LY1 antibodies were generated by immunizing rabbits with a synthetic peptide representing one of three portions of the ORF1 protein (jelly roll residues 46-58: KIKRLNIVEWQPK, spike domain residues 485-502: SPSDTHEPDEEDQNRWYP, C-terminal domain residues 635-672: SEEEEESNLFERLLRQRTKQLQLKRRIIQTLKDLQKLE) conjugated to a carrier protein by an engineered N-terminal cysteine. Rabbits were immunized twice, and polyclonal antibodies were purified from bleeds using protein A purification (Custom Antibody Production, Life Technologies corporation). Western blot analysis was performed using NUPAGE 4-12% gels (ThermoFisher) transferred to nitrocellulose membranes using the Transblot Turbo system (BioRad). Membranes were blocked with blocking buffer (Licor), probed ∼16 hours with primary antibody, and detected using anti-rabbit IRDye infrared secondary antibodies and imaging system (Licor).

### Construct design, cell culture and protein expression/purification

The LY1 ORF2 and ORF1 sequences^13^ were codon optimized for insect cells and with different ORF1 construct length variations (full-length ORF1, ΔARM with a deletion of residue 2-45, and ΔARM/ΔC-term with deletions of residues 2-45 and 552-672). The ORF2 and ORF1 constructs were cloned into the pFastBac Dual plasmid which was used to generate baculoviruses (Genscript and Medigen). To express the ORF1 proteins, Sf9 cells (Gibco™ 11496015) were infected by baculovirus with multiplicity of infection = 1 and the cells were cultured for three days at 27°C and harvested by centrifugation. The cells were lysed by treatment with 0.01% Triton X-100 (Sigma-Aldrich 11332481001), subjected to micro-fluidization, and treated with protease inhibitors (Thermo Scientific Halt Protease Inhibitor Cocktail, PI78438) and DNase (Benzonase^®^; Sigma). Cell lysate was subsequently purified using HiTrap Heparin affinity chromatography (Cytiva) followed by size-exclusion chromatography (HiPrep 16/60 Sephacryl S-500 HR; Cytiva).

### Negative-stained EM data collection and analysis

We used Jeol 1200EX transmission electron microscope for screening different TTMV-LY1 constructs. 10 μl of sample was blotted on 400 mesh carbon support film (cf400-cu, EMS) for 30 seconds. After washing by ddH2O for 30 seconds, the grid was stained by 0.75% of uranyl formate (UF) for 10 seconds before loading on the scope. LY1 ΔARM was further imaged at NanoImaging Service. 3 μl of 0.12 mg/ml was blotted to a continuous carbon grid and stained by 1% UF. Negative-stained electron microscopy was performed using a Thermo Fisher Scientific (Hillsboro, Oregon) Glacios Cryo Transmission Electron Microscope (cryo-TEM) operated at 200 kV and equipped with a TFS CETA-D 4×4 CMOS camera and a Falcon 4 direct electron detector.

### Cryo-EM data collection and data analysis and molecular refinement

For the grid preparation, 3 μl of 0.3 mg/ml VLP sample was applied to a 1.2×1.3 graphene oxide grid. A total of 11,083 micrographs were collected from Glacios cryo-TEM (Thermo Fisher Scientific) operated at 200 kV with a Falcon 4 direct electron detector, at a nominal defocus range of -1.0 – -2.5 μm and accumulated dose of 19.59 e^-^/Å^22^ for a total of 15 frames in 3 minutes. The pixel size was 0.923 Å, with the magnification 150000x. Automated data-collection was carried out by Leginon^22^ software.

All micrographs were motion-corrected by Relion-4.0^23,24,25^ implemented MotionCor2^26^, and the contrast transfer function (CTF) parameters were estimated by Gctf^27^. With manually picked particles from 20 micrographs to train the network, SPHIRE-crYOLO^28^ automatically picked 58,391 particles along with the PhosaurusNet network. All particles were extracted by Relion-4.0 and rescaled to 2-fold (pixel size 1.846 Å), followed by subsequence 2D classifications with 350 Å mask diameter to remove any junk particles. Two iterations of 2D classifications resulted in 11,185 particles, which were merged and reextracted to generate a de novo 3D initial model by Relion-4.0 with I1 symmetry. Notably, several similar initial models without symmetry imposed were obtained by Relion-4.0 and cisTEM^29^. To obtain a better classification result, all particles were first subjected to a Refine3D with initial angular sampling 3.7° and local angular search 0.9° per step. The alignment parameters of each particle were transferred to a 3D classification with angular sampling interval 0.9° and local angular search 5° per step. 3D classification of the entire particle set attributed most of the particles into a single class. After CtfRefine and Bayesian polishing, the post-processing results in 3.98 Å resolution under the gold standard (with FSC=0.143).

The initial anellovirus TTMV-LY2^13^ capsid monomer structure was predicted by TrRosetta^30^, and TTMV-LY1 structure was further predicted by RosettaCM^31^. Structural refinement was performed by Rosetta^32^ and Phenix^33^, and fine-adjusted by COOT^34^. JA 20 and SAFIA structures are predicted by Alphafold.^35^

### Sequence alignments

Sequences of select anellovirus ORF1s were taken from Genebank and aligned using Clustal Omega. ORF1 sequences used in the alignment are indicated by their accession numbers in Extended Data Figure 4, except for the four sequences also shown in Figures 2 and 3, who are indicated with abbreviated names for clarity. The accession numbers of these sequences are LY1 (YP 007518450.1), LY2 (AGG91484.1), SAfiA (MN779270.1) and JA20 (AF122914.3). The consensus sequence is a composite of residues conserved (identity greater than 30%) or similar residues (greater than 50%; hydrophobic and bulky aromatic (Tyr) residues indicated with ϕ and hydrophilic and basic residues indicated with γ) Alignments and consensus sequences are shown in Figures 2, 3 and Extended Data Figure 4.

### Circular dichroism

To determine the secondary structure of the LY1 C-terminal domain, the peptides used to generate C-terminal antibodies (residues 635-672) were analyzed at 25.8 μM in PBS on a Jasco J-815 circular dichroism spectropolarimeter using 2 mm path length cell at ambient temperature. Each data set is the average of three consecutive scans. The secondary structure was determined by using CDPro software package^36^, compared with a reference set containing 56 proteins (IBasis=10). The final secondary structure fractions were averaged over the results from three programs (SELCON3, CDSSTR, CONTINLL) in CDPro.

## Acknowledgements

We thank Dr. Joseph Che-Yen Wang for manuscript discussions and suggestions, and Maria Ericsson and the team at Harvard Medical School Electron Microscopy facility for their technical support. Cryo-EM data were collected at NanoImaging Services under the leadership of Dr. Giovanna Scapin and Dr. Phat Dip. Baculovirus was generated by Dr. Peter Pushko at Medigen.

## Author Contributions

S.-H. L. purified samples, conducted negative EM, and processed the data with structural refinement. N.C. initiated the molecular model. Y.Z. and N.C. performed circular dichroism measurement. N.M.A., H.R., S.I., and L.Z. performed ORF1 purification and construct screening. S.D., R.H., T.O. and Y.C. oversaw the project. K.S. designed the ORF1 constructs and oversaw the project. S.-H. L., S.D., and K.S. wrote the manuscript.

## Competing Interest Declaration

All authors are employed and hold equity interests in Ring Therapeutics. S.-H. L., N.C., S.D., and K.S. are the inventors on a patent application related to this work.

## Extended Data Tables

**Extended Data Table 1.**
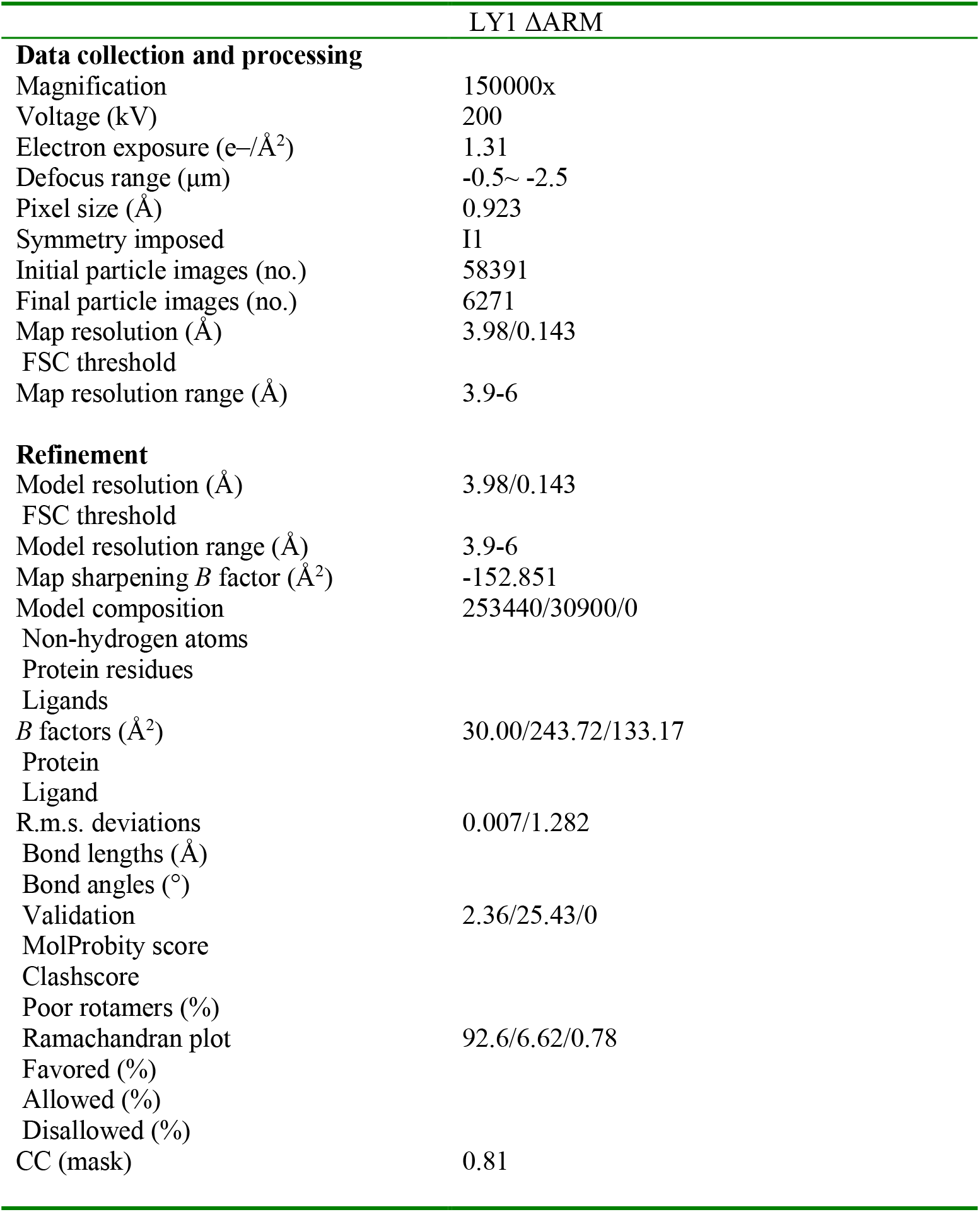
Cryo-EM data collection, refinement and validation statistics of TTMV-LY1 ΔARM.

## Extended Data Figures

**Extended Data Fig. 1.**
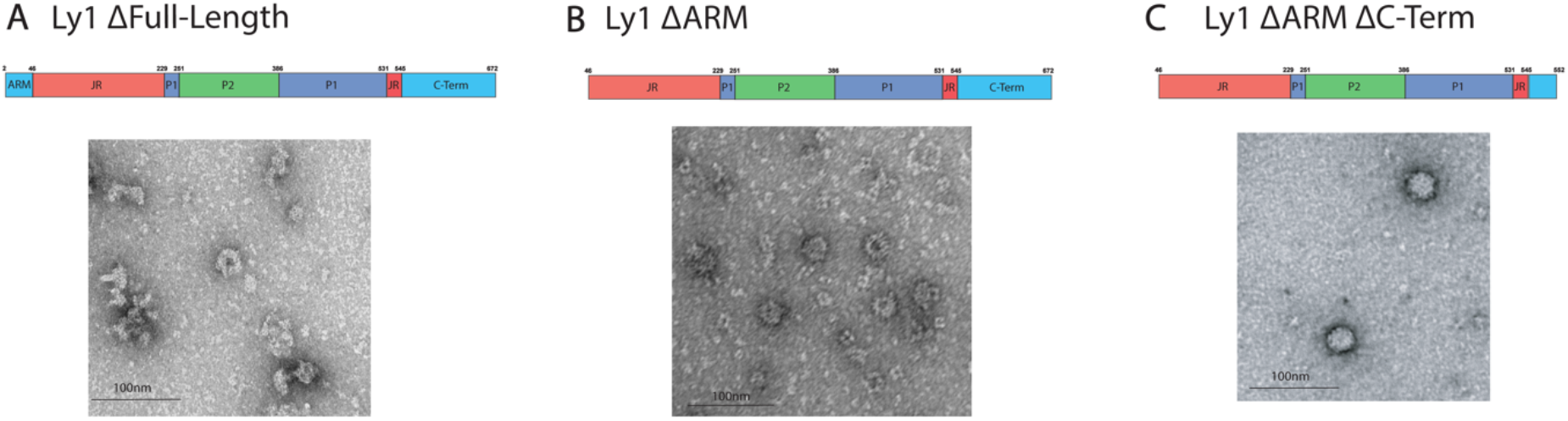
Comparison of TTMV-LY1 constructs by negative-stained electron microscopy. **A.** Full-length LY1 shows highly heterogeneous particles. **B**. LY1 ΔARM virus-like particles have a more symmetrical appearance after C-terminal proteolysis. **C**. Genetic truncation of the C-terminal (Δ552-672) preserves a symmetrically structured particle. Scale bar = 100 nm.

**Extended Data Fig. 2.**
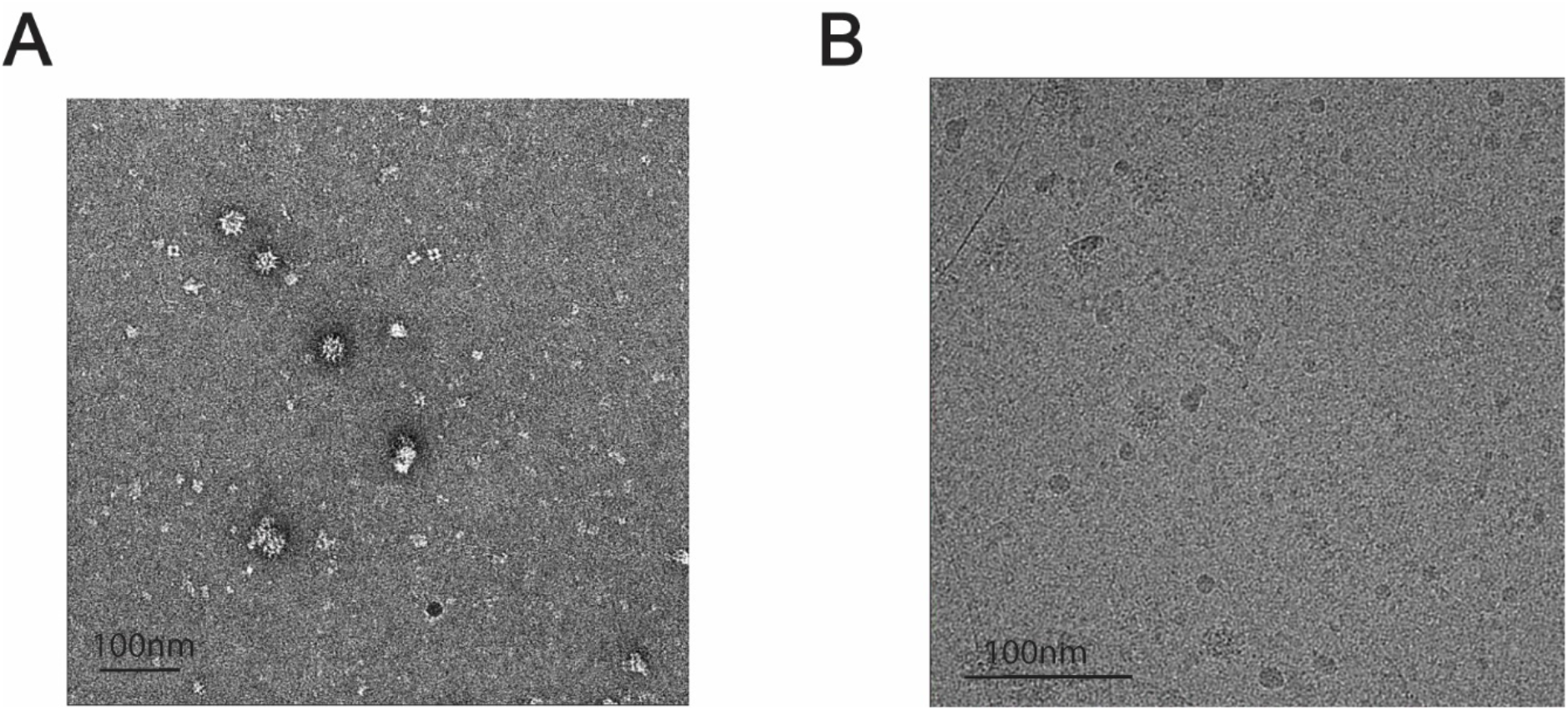
Representative micrographs of TTMV-LY1 ΔARM. **A** and **B** are representative negative-stained and cryo-EM micrographs for LY1 ΔARM, respectively.

**Extended Data Fig. 3.**
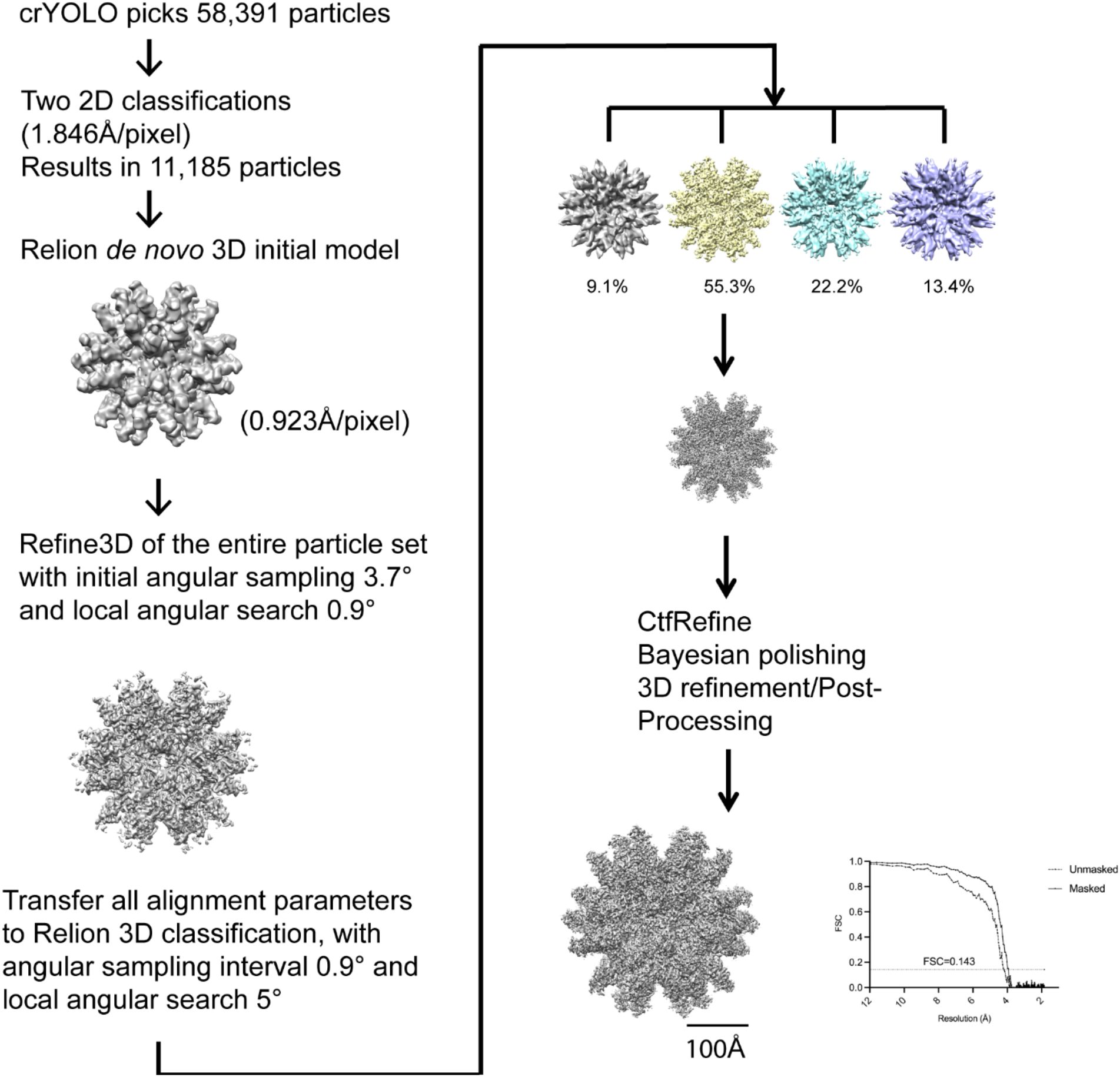
Data processing procedure of the LY1 ΔARM cryogenic electron microscopy (cryo-EM) reconstruction. In short, crYOLO picks 58,391 particles from 11,083 micrographs. Several rounds of 2D classification result in 11,185 particles. After Relion de novo initial model reconstruction, Relion 3D refinement was implemented to obtain the orientation parameters. All particles with parameters are fed in a 3D classification. The class with the most abundant particle population results in 3.98 Å resolution.

**Extended Data Fig. 4.**
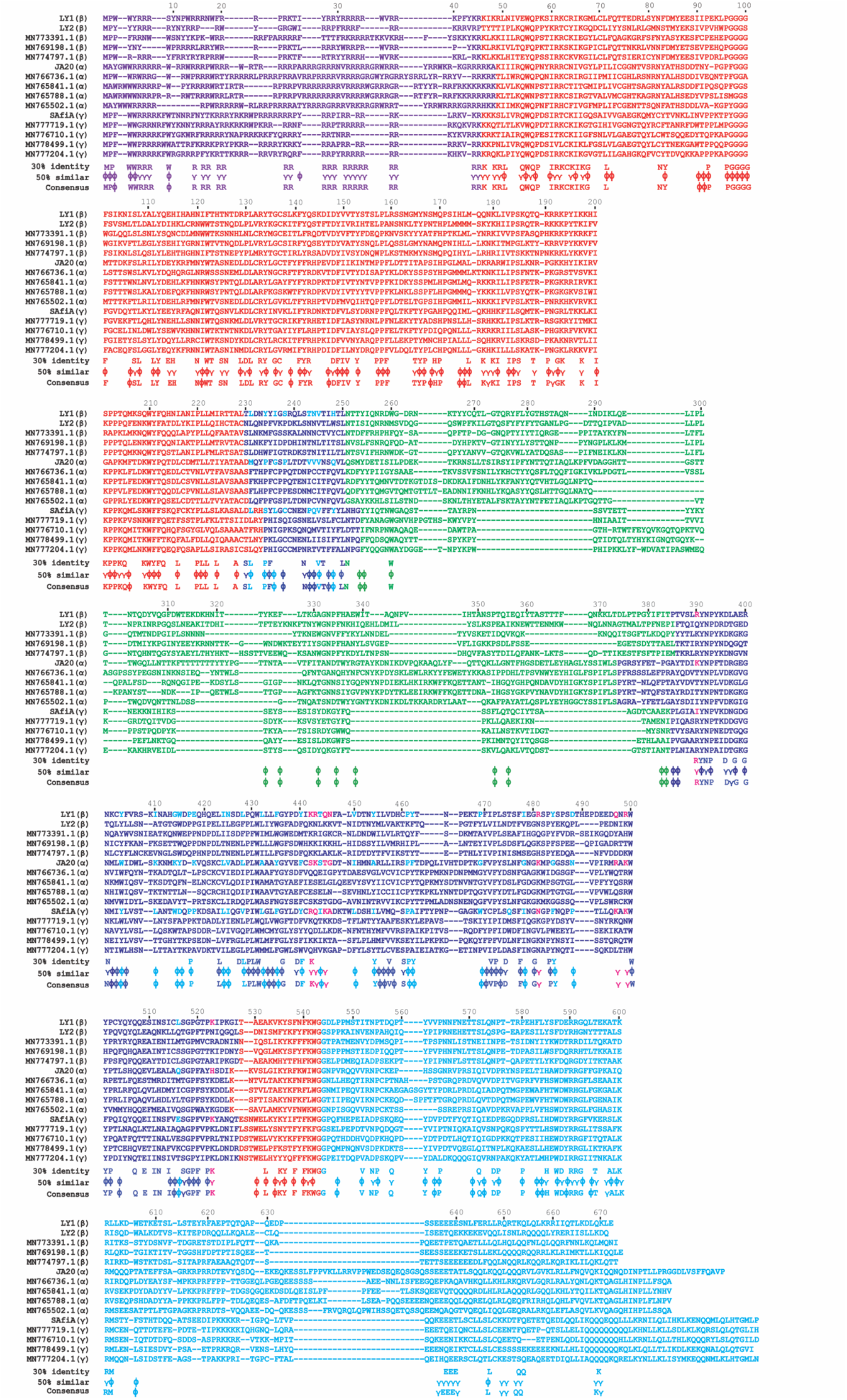
Sequence alignment of 15 published anelloviruses within different genera indicated in parentheses. The residues are colored by domain as in Figure 1. Below the alignment, conserved residues (>30%) or similar (>50%; hydrophobic and Tyr with ø and hydrophilic with γ) are shown below the alignment. Surface exposed hydrophobic (cyan) and hydrophilic (magenta) are colored as in Figure 3. A consensus sequence combining conserved and similar residues is also shown.

**Extended Data Fig. 5.**
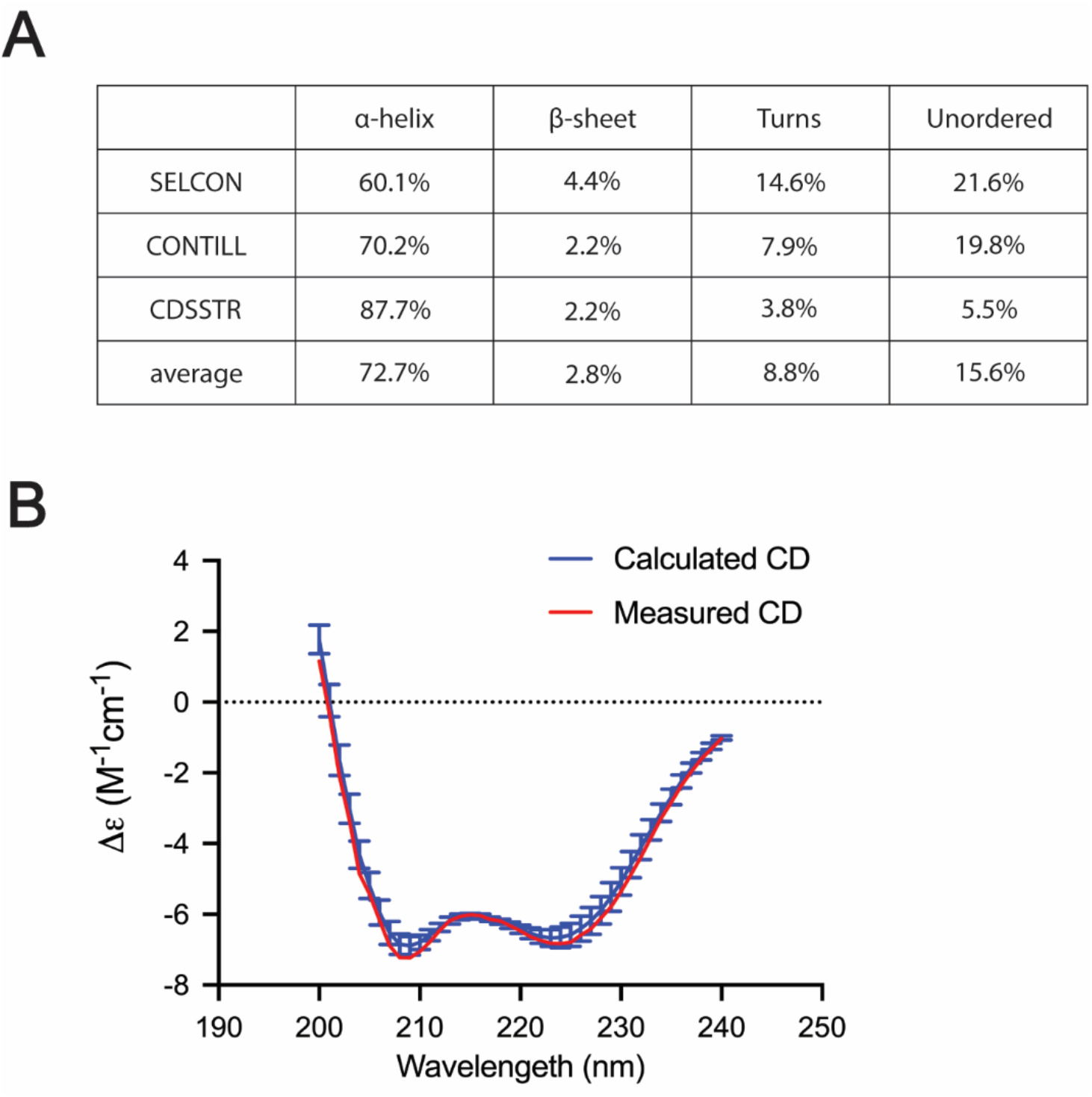
Circular dichroism (CD) of the LY1 C-terminal peptide. **A.** Averages of secondary structure fractions estimated by different packages of CDPro. α-helix dominates the secondary structure assignment from the CD spectrum. **B**. An experimental spectrum of the C-terminal peptide (shown in red) overlaid with the calculated and averaged reference set spectra (shown in blue).

## Notes

### Competing Interest Statement

All authors are or were employed and hold equity interests in Ring Therapeutics. S.-H. L., N.C., S.D., and K.S. are the inventors on a patent application related to this work.

